# Single-cell sequencing of trophoblasts in preeclampsia and chemical hypoxia in BeWo b30 cells reveals EBI3, COL17A1, miR-27a-5p and miR-193b-5p as hypoxia-response markers

**DOI:** 10.1101/2025.09.30.679431

**Authors:** Evgeny Knyazev, Timur Kulagin, Ivan Antipenko, Alexander Tonevitsky

**Affiliations:** Faculty of Biology and Biotechnology, HSE University, Moscow, Russia; Laboratory of Microfluidic Technologies for Biomedicine, Shemyakin-Ovchinnikov Institute of Bioorganic Chemistry of the Russian Academy of Sciences, Moscow, Russia

**Author notes:** Corresponding author, Faculty of Biology and Biotechnology, HSE University, 33k4 Profsoyuznaya Street, Moscow, Russia *E-mail address* (E. Knyazev).

**Keywords:** Preeclampsia, trophoblast, single-cell RNA sequencing, hypoxia, microRNA

## Abstract

**Background:** Preeclampsia (PE) complicates 2–8% of pregnancies and is marked by placental hypoxia and HIF-pathway activation, especially in early-onset PE (eoPE). Integrating patient tissue analyses with experimental models may reveal common molecular markers of trophoblast hypoxic response.

**Methods:** We analyzed scRNA-seq data from 10 eoPE, 7 late-onset PE (loPE), and corresponding control placentas, identifying villous cytotrophoblast (VCT), syncytiotrophoblast (SCT), and extravillous trophoblast (EVT) subpopulations. BeWo b30 cells were treated for 24 h with CoCl_2_ (300 µM) or an oxyquinoline derivative (OD, 5 µM) to induce hypoxia. RNA and small RNA sequencing quantified mRNA and microRNA changes. PROGENy inferred pathway activities.

**Results:** Single-cell analysis revealed highest hypoxia pathway activation in eoPE, with EVT showing maximum activity among trophoblast populations. Nine genes were upregulated across all trophoblast types in eoPE: *EBI3, CST6, FN1, RFK, COL17A1, LDHA, PKP2, RPS4Y1*, and *RPS26. In vitro*, OD induced more specific hypoxia responses than CoCl_2_, with 1,284 versus 3,032 differentially expressed genes respectively. Critically, *EBI3, FN1*, and *COL17A1* showed concordant upregulation in both placental tissue and OD-treated cells, while CoCl_2_ treatment produced opposite expression patterns. MicroRNA analysis identified hsa-miR-27a-5p and hsa-miR-193b-5p as consistently elevated in both experimental conditions and previously reported in PE placental vesicles. We also identified isoforms of hsa-miR-9-5p and hsa-miR-92b-3p as hypoxia-associated in trophoblast.

**Conclusions:** *EBI3, COL17A1*, hsa-miR-27a-5p, and hsa-miR-193b-5p emerge as trophoblast hypoxia markers in PE. Oxyquinoline derivatives offer a more physiologically relevant *in vitro* hypoxia model than CoCl_2_. This integrated approach advances understanding of PE pathophysiology and suggests candidate therapeutic targets.

## 1. Introduction

Preeclampsia (PE) is a serious pregnancy complication, affecting 2–8% of pregnancies worldwide and representing a leading cause of maternal and perinatal morbidity and mortality [1]. This complex, multisystem disorder originates from defective placentation in early gestation, characterised by inadequate extravillous trophoblast invasion of the maternal decidua and impaired spiral-artery remodelling [2]. These abnormalities precipitate placental ischaemia and hypoxia, processes central to PE pathophysiology [3].

Understanding the molecular pathology of PE is challenging due to its clinical heterogeneity, which encompasses subtypes with distinct presentations and outcome. Early-onset PE (eoPE; onset before 34 weeks’ gestation) is characterized by marked placental dysfunction and activation of hypoxia-responsive pathways, notably those mediated by HIF-1α [4]. In contrast, late-onset PE (loPE) has a less clear association with placental abnormalities, implying that alternative pathophysiological processes may predominate [4].

*In vitro* trophoblast models, including BeWo b30 cells, are widely employed to investigate the molecular underpinnings of PE. These cells are treated with hypoxia-mimetic agents such as cobalt(II) chloride (CoCl_2_) and oxyquinoline derivatives (ODs), which inhibit HIF-prolyl hydroxylases [5–9]. However, the precise mechanisms by which these mimetics act differ.

CoCl_2_ is commonly used to mimic hypoxia *in vitro*, but its actions are pleiotropic. The primary mechanism by which CoCl_2_ mimics hypoxia is thought to be the substitution of Fe^2+^ with Co^2+^ in the catalytic site of HIF prolyl hydroxylases (PHDs), thereby inhibiting their activity and stabilizing HIF-1α [10]. Additional proposed mechanisms include oxidation of ascorbate cofactors essential for PHD function, direct binding to HIF-1α that blocks interaction with the von Hippel–Lindau protein (pVHL), and inhibition of the asparaginyl hydroxylase FIH [10]. However, CoCl_2_ also elicits significant off-target effects: it generates reactive oxygen species and depletes intracellular glutathione, impairs mitochondrial membrane potential and ATP production, activates p53-dependent apoptotic pathways, induces DNA damage independently of hypoxia, and perturbs iron homeostasis by competing for metal transport proteins [10].

As a more specific hypoxia model, ODs have been developed as selective PHD inhibitors. ODs bind directly to the Fe^2+^ ion in the PHD active site without perturbing other iron-dependent enzymes [11]. Substituents at the 7-position of the oxyquinoline ring mimic the HIF-1α oxygen-dependent degradation domain that engages the PHD active site, conferring high binding specificity. Consequently, ODs selectively block HIF-1α hydroxylation, leading to its stabilization and activation of hypoxia-responsive pathways without overt off-target toxicity [12,13]. Despite widespread use of CoCl_2_ and ODs to model trophoblast hypoxia in PE research, it remains unclear how faithfully these approaches recapitulate the complex gene-expression profiles characteristic of PE placental tissue.

Whole-tissue RNA sequencing (RNA-seq) of placental lysates has long underpinned PE research, with differentially expressed genes identified by comparison to normotensive controls. Chemical induction of hypoxia in trophoblast cell lines offers a controlled *in vitro* model of this pathology. However, bulk RNA-seq and single-cell RNA sequencing (scRNA-seq) of distinct trophoblast subsets often yield divergent signatures, reflecting the cellular heterogeneity of placental tissue [14]. Advances in scRNA-seq now enable high-resolution interrogation of molecular alterations within villous cytotrophoblasts (VCT), syncytiotrophoblasts (SCT), and extravillous trophoblasts (EVT)—each with specialized roles in placentation—providing unprecedented insight into subtype-specific gene-expression patterns in PE [15].

In this study, we examined gene-expression alterations in three trophoblast subpopulations in PE using scRNA-seq data and compared these profiles with those of BeWo b30 cells exposed to CoCl_2_ or OD. We also assessed the expression of placental extracellular vesicle markers in PE versus normotensive pregnancies. Our aim was to delineate and contrast the molecular responses of primary trophoblast subsets and hypoxia-treated cell lines, thereby identifying potential biomarkers of hypoxic perturbation in PE.

## 2. Materials and methods

### 2.1. Single-cell RNA-seq data acquisition

Single-cell RNA sequencing data were obtained from Admati et al. (2023) [16], generated on the 10x Genomics Chromium platform. The authors made available a final dataset consisting of a genes × cells UMI count matrix together with full cell metadata for 10 eoPE cases, 7 loPE cases, and corresponding gestational age-matched normotensive control placentas (three matched to eoPE and six to loPE cases). Count data and metadata were imported into Python (version 3.13.7) as an AnnData object using the scanpy package (version 1.11.4).

### 2.2. Data processing and quality control

Raw UMI counts underwent initial filtering in scanpy: cells with <200 genes, genes detected in <3 cells, or >10% mitochondrial counts were removed. Doublets were identified and excluded using scrublet. Filtered counts were normalized per cell to 10,000 total counts with sc.pp.normalize_total(), log-transformed with sc.pp.log1p(), and highly variable genes selected with sc.pp.highly_variable_genes(). Dimensional reduction and clustering were performed via PCA (sc.tl.pca()), neighbor graph construction (sc.pp.neighbors()), and UMAP embedding (sc.tl.umap()). Detailed subclusters defined by [16] were aggregated into three trophoblast categories (VCT, SCT, EVT) for comparative analysis.

### 2.3. Differential gene expression in single-cell RNA-seq data and pathway analysis

Differential expression between PE and control samples was computed on the log-normalized data using sc.tl.rank_genes_groups() (Wilcoxon rank-sum test with Benjamini–Hochberg correction). Genes with |log_2_ fold change| > 0.585 and adjusted p < 0.05 were deemed significant. Overlap among gene sets was visualized using jvenn [17], and genes consistently upregulated in eoPE across all trophoblast subtypes were designated common PE markers. Pathway activities were inferred on cluster-averaged expression profiles using PROGENy (v2.1.1) to compute scores for fourteen signaling pathways [18].

### 2.4. Statistical analysis and visualization of single-cell RNA-seq data

All analyses were conducted in Python using scanpy and scverse packages. Non-parametric tests determined statistical significance (adjusted p < 0.05). Data visualization, including UMAP plots and violin plots of gene expression, was generated with scanpy’s plotting functions and matplotlib. All workflows utilized AnnData objects for efficient handling of large single-cell datasets.

### 2.5. Hypoxia induction in a trophoblast cell model

BeWo b30 choriocarcinoma cells—a well-established trophoblast model—were kindly provided by Prof. Dr. Christiane Albrecht (University of Bern, Switzerland) with permission from Prof. Dr. Alan Schwartz (Washington University in St. Louis, USA). Cells were cultured in DMEM supplemented with 10% fetal bovine serum, 1% non-essential amino acids, and 1% penicillin–streptomycin (Gibco, USA) as described previously [19]. At ~80% confluence, cells were treated for 24 h with either 300 µM cobalt(II) chloride (Sigma-Aldrich, USA) or 5 µM oxyquinoline derivative 4896-3212 (ChemRar, Russia) in 0.05% DMSO to induce HIF-1α stabilization; controls received medium with 0.05% DMSO. After treatment, cells were lysed in QIAzol, and total RNA was extracted using the miRNeasy Mini Kit with on-column DNase digestion (QIAGEN, Germany).

### 2.6. RNA sequencing and analysis for cell model

mRNA and miRNA libraries were prepared using the Illumina Stranded mRNA Library Prep Kit and the NEBNext Small RNA Library Prep Kit, respectively. Sequencing was performed on an Illumina NextSeq 550 platform (75-bp single-end reads for mRNA; 50-bp reads for miRNA).

Raw reads were quality-trimmed and aligned to the human genome (GRCh38). miRNA isoforms (isomiRs) were detected and quantified with IsomiRmap. Library sizes were normalized by trimmed mean of M-values (TMM) normalization using edgeR v4.6.3 [20]. Gene expression was quantified as log_2_-transformed fragments per kilobase of transcript per million mapped reads (FPKM) for mRNA and reads per million mapped reads (RPM) for miRNA. Low-abundance transcripts—defined as the lowest 25% of mRNAs and the lowest 50% of miRNAs by median expression—were filtered out [21,22] to account for the skewed distribution of miRNA abundance [23].

Differential expression analysis was conducted with DESeq2 v1.48.2, applying Benjamini–Hochberg false discovery rate (FDR) correction. For mRNA-seq, FC shrinkage was performed using apeglm, with s-values <0.005 and an absolute FC ≥2 considered significant. For miRNA-seq, significance thresholds were FDR <0.05 and absolute FC ≥1.5. Functional annotation of common markers was performed using DAVID to identify enriched annotation clusters [24]. The obtained RNA-seq results have been deposited in the Gene Expression Omnibus (GEO) under accession number GSE308908.

### 2.7. Real-time PCR analysis

To validate differential gene expression, quantitative PCR (qPCR) was performed for selected genes. RNA concentration and purity were assessed using a NanoDrop spectrophotometer, and RNA integrity was confirmed by agarose gel electrophoresis. First-strand cDNA synthesis was performed using 1.5 μg of total RNA with the MMLV RT kit (Evrogen, Russia) according to the manufacturer’s instructions. The cDNA was diluted 1:20 in nuclease-free water prior to qPCR analysis.

qPCR reactions were conducted using the 5X qPCRmix-HS SYBR kit (Evrogen, Russia) in a total volume of 25 μL. Each reaction contained 5 μL of 5X qPCRmix-HS SYBR master mix, 1.25 μL each of forward and reverse primers (5 μM stock), 2 μL of diluted cDNA template, and 15.5 μL of nuclease-free water. The thermal cycling conditions consisted of an initial denaturation at 95°C for 5 minutes, followed by 40 cycles of denaturation (95°C, 20 seconds), annealing (64°C, 10 seconds), and extension (72°C, 15 seconds). Melting curve analysis was performed to verify amplification specificity.

Gene-specific primers were designed using NCBI Primer-BLAST (Table 1) and synthesized by Evrogen (Russia). *ACTB* served as the reference gene for normalization. All reactions were performed in technical triplicates, and biological replicates (n = 3) were analyzed for each experimental condition. Relative gene expression was calculated using the 2^−ΔΔCt^ method, with fold-change values normalized to control samples. Statistical analysis was performed using Student’s t-test, with p < 0.05 considered statistically significant.

**Table 1.**
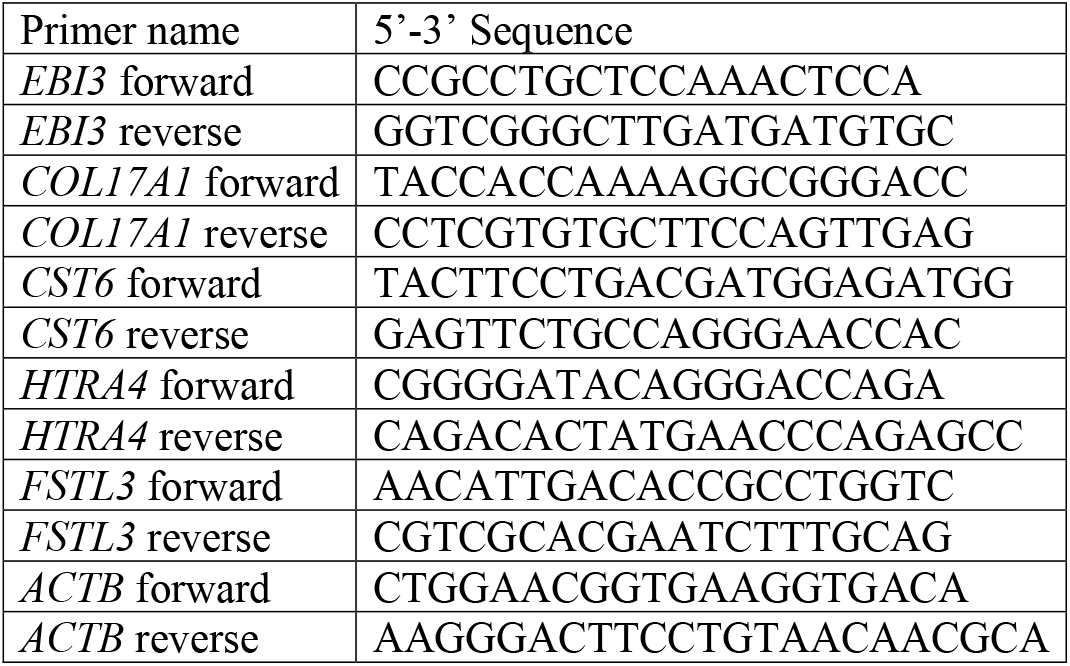
qPCR primer sequences.

## 3. Results

### 3.1. Analysis of single-cell sequencing data

#### 3.1.1. Trophoblast subpopulation delineation reveals skewed cluster distributions in early- and late-onset preeclampsia

We analyzed a publicly available scRNA-seq dataset [16] to delineate trophoblast subpopulations in PE. The original study profiled ten eoPE placentas (diagnosed before 34 weeks’ gestation) alongside three gestationally matched controls, and seven loPE placentas with six matched normotensive controls. They defined the following trophoblast subsets: proliferating VCT (VCT_PROLIFERATING) and two postmitotic VCT clusters (VCT_1 и VCT_2); small VCT subset with an interferon-response signature (VCT_IFN); two EVT clusters, EVT_1 (high *PAPPA* and *PRG2* expression) and EVT_2 (*TNNI2* expression); classic SCT cluster with *CYP19A1* and *CGA* expression (SCT) and small placenta specific gene (PSG)-enriched SCT subset additionally expressing *PSG1, PSG3, PSG9, PSG11*, and *KISS1* (SCT_PSG). We reclustered these populations for eoPE and matched controls (Figure 1A), aggregated EVT, SCT, and VCT clusters for combined analysis (Figure 1B), and compared the proportional distribution of each cluster between PE and control samples (Figure 1C). The same workflow was applied to the loPE cohort (Figure 1D–F).

**Fig. 1.**
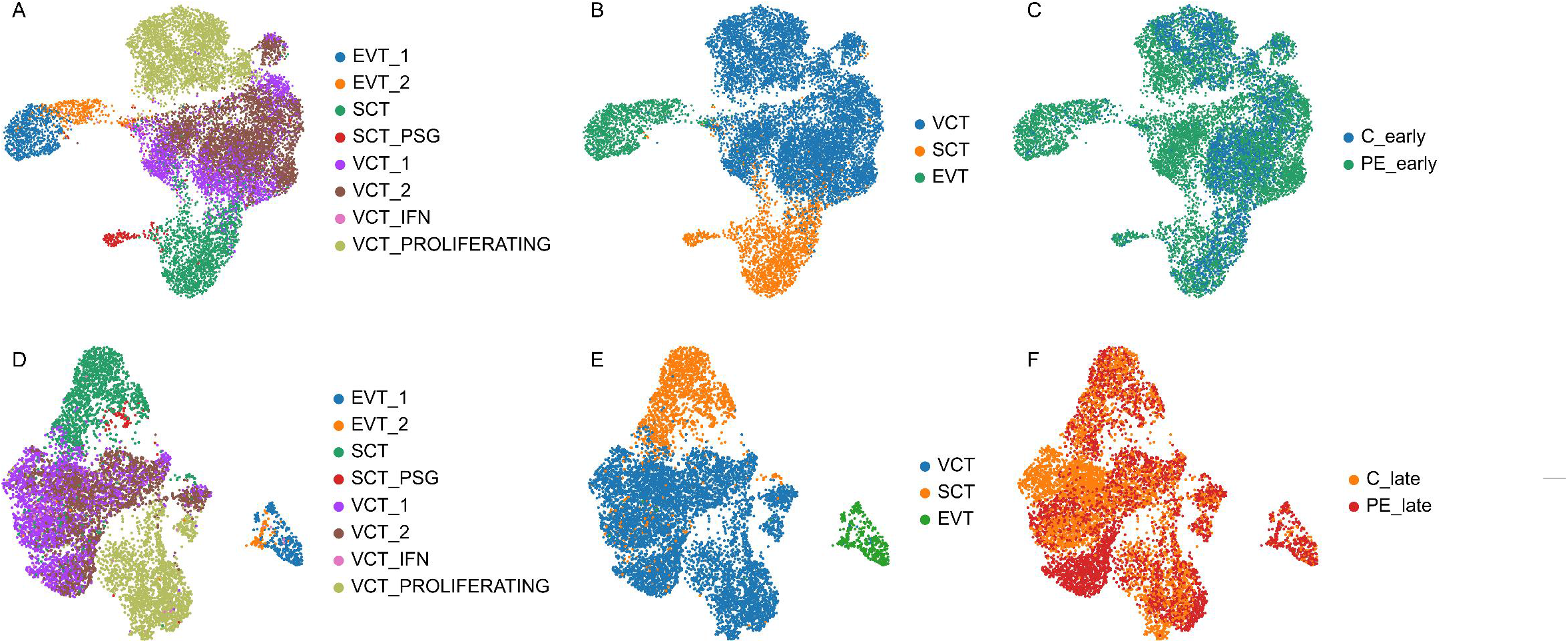
UMAP clustering of trophoblast subpopulations based on single-cell RNA-seq data. **A–C**. Early-onset PE (eoPE) and matched controls. **D–F**. Late-onset PE (loPE) and matched controls. **A, D**. Annotation of trophoblast subclusters as defined by [16]. **B, E**. Aggregated trophoblast clusters. **C, F**. Relative distribution of cells between preeclampsia (PE) and control (C) samples. EVT_1, extravillous trophoblasts (EVT) with high *PAPPA* and *PRG2* expression; EVT_2, EVT with *TNNI2* expression; SCT, classic syncytiotrophoblast (SCT) fraction marked by *CYP19A1* and *CGA*; SCT_PSG, minor SCT subset additionally expressing *PSG1, PSG3, PSG9, PSG11*, and *KISS1*; VCT_ PROLIFERATING, proliferating villous cytotrophoblast (VCT); VCT_1 and VCT_2, postmitotic VCT clusters; VCT_IFN, VCT subset exhibiting active interferon response; C_early, controls for eoPE (delivery <34 weeks); PE_early, eoPE samples; C_late, controls for loPE (delivery ≥34 weeks); PE_late, loPE samples.

Cell counts for each trophoblast cluster and subcluster are summarized in Table 2. Although the early controls yielded the fewest total cells (3,442), followed by late controls and loPE samples (4,265 and 4,299, respectively), and eoPE samples the most (9,867), the per-sample averages were 987 cells for eoPE and 1,147 for early controls, and 614 cells for loPE and 711 for late controls. Notably, the VCT:SCT ratio ranged from 3.8:1 in loPE to 5:1 in eoPE, likely reflecting underrepresentation of SCT in scRNA-seq due to loss of large syncytial structures during tissue dissociation and cell capture.

**Table 2.** Cell counts across trophoblast clusters in single-cell RNA sequencing data.

| Condition | Trophoblast subclusters as defined by [16] |  |  |  |  |  |  |  | Aggregated trophoblast clusters |  |  |
| --- | --- | --- | --- | --- | --- | --- | --- | --- | --- | --- | --- |
|  | VCT_1 | VCT_2 | VCT_IFN | VCT_PROLIFERATING | SCT | SCT_PSG | EVT_1 | EVT_2 | VCT | SCT | EVT |
| C_early | 603 | 1203 | 27 | 895 | 632 | 26 | 31 | 25 | 2728 | 658 | 56 |
| C_late | 1194 | 1307 | 24 | 915 | 696 | 28 | 81 | 20 | 3440 | 724 | 101 |
| PE_early | 1947 | 2984 | 79 | 2229 | 1272 | 174 | 797 | 385 | 7239 | 1446 | 1182 |
| PE_late | 1037 | 1100 | 64 | 977 | 793 | 48 | 217 | 63 | 3178 | 841 | 280 |

VCT\_1 and VCT\_2, postmitotic villous cytotrophoblast (VCT) clusters; VCT\_IFN, VCT subset exhibiting active interferon response; VCT\_PROLIFERATING, proliferating VCT; SCT, classic syncytiotrophoblast (SCT) fraction marked by *CYP19A1* and *CGA*; SCT\_PSG, minor SCT subset additionally expressing *PSG1*, *PSG3*, *PSG9*, *PSG11*, and *KISS1*; EVT\_1, extravillous trophoblasts (EVT) with high *PAPPA* and *PRG2* expression; EVT\_2, EVT with *TNNI2* expression; C\_early, controls for early-onset preeclampsia (delivery <34 weeks); C\_late, controls for late-onset preeclampsia (delivery ≥34 weeks); PE\_early, early-onset preeclampsia samples; PE\_late, late-onset preeclampsia samples.

#### 3.1.2. Signaling pathway analysis reveals elevated hypoxia and inflammatory signaling in EVT and SCT subpopulations

To assess signaling pathway activity in trophoblast clusters, we utilized both the detailed subclusters defined by [16] and aggregated clusters. First, we combined all subclusters within eoPE and loPE cohorts and their respective controls to generate integrated pathway activity profiles using PROGENy—a tool that infers activity of 14 key oncogenic and signaling pathways from target gene expression [18]. The results (Figure 2) revealed that hypoxia pathway activity was highest in eoPE samples, intermediate in loPE samples, and lowest in controls. Additionally, eoPE exhibited the strongest activation of NFκB, PI3K, TGF-β, TNF-α, and TRAIL pathways, suggesting heightened inflammatory and apoptotic processes.

**Fig. 2.**
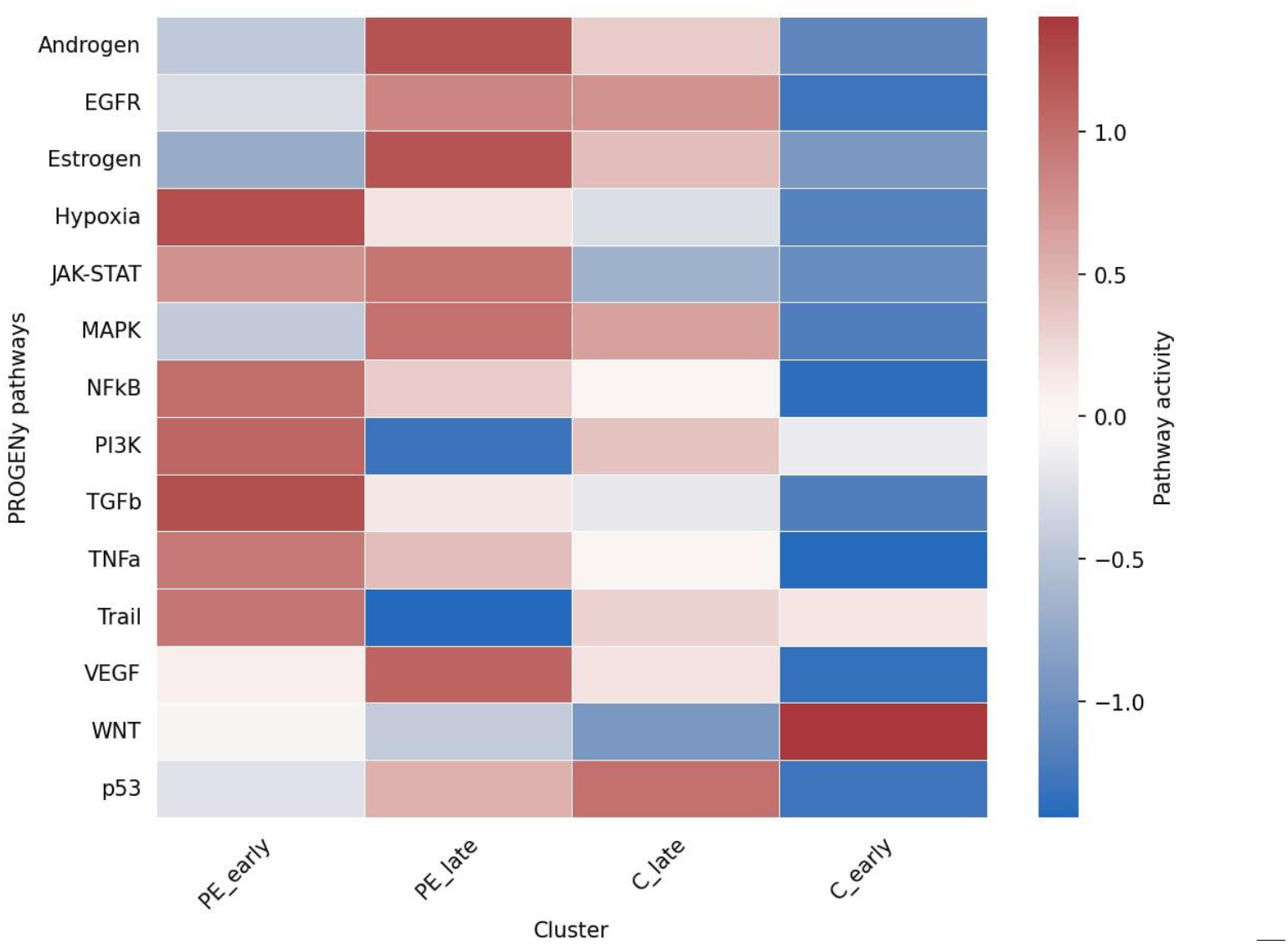
PROGENy-inferred signaling pathway activities in aggregated trophoblast clusters. PE_early, early-onset preeclampsia (eoPE) samples; PE_late, late-onset preeclampsia (loPE) samples; C_late, controls for loPE (delivery ≥34 weeks); C_early, controls for eoPE (delivery <34 weeks).

We next evaluated PROGENy-inferred pathway activities within the specific trophoblast subclusters defined by [16], irrespective of disease status (Figure 3). Hypoxia signaling was most pronounced in EVT, consistent with their distance from maternal blood vessels. The SCT_PSG subset—considered the most representative syncytiotrophoblast fraction given loss of large syncytial cells during dissociation— exhibited the next highest hypoxia activity. Both EVT and SCT_PSG also showed elevated NFκB, PI3K, TGF-β, TNF-α, and TRAIL pathway activities, pathways known to be activated in PE.

**Fig. 3.**
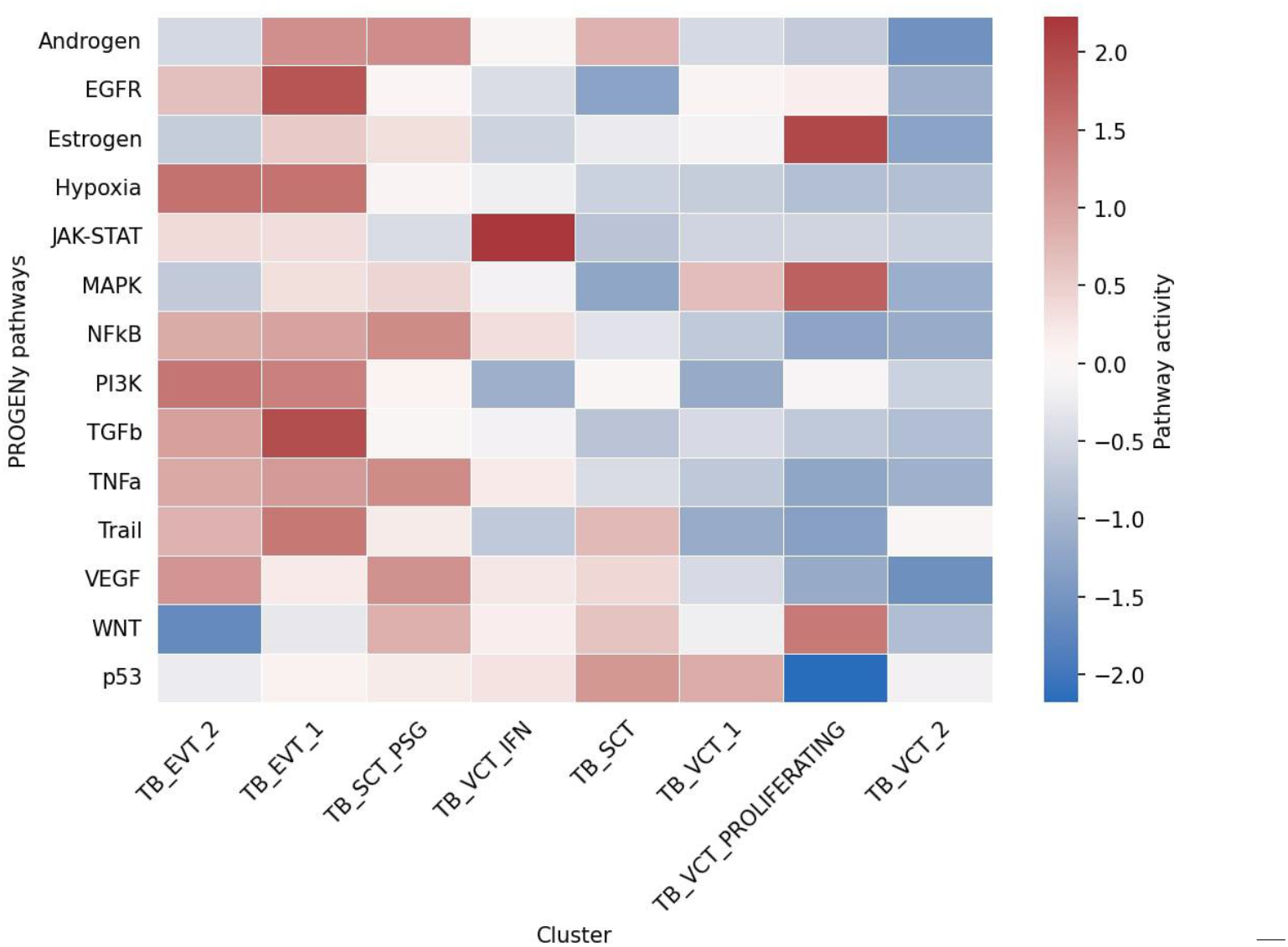
PROGENy-inferred signaling pathway activities across trophoblast subclusters from scRNA-seq data [16]. TB, trophoblast; TB_EVT_1, extravillous trophoblast (EVT) with high *PAPPA* and *PRG2* expression; TB_EVT_2, EVT with *TNNI2* expression; TB_SCT, classic syncytiotrophoblast (SCT) fraction marked by *CYP19A1* and *CGA*; TB_SCT_PSG, minor SCT subset additionally expressing *PSG1, PSG3, PSG9, PSG11*, and *KISS1*; TB_VCT_ PROLIFERATING, proliferating villous cytotrophoblast (VCT); TB_VCT_1 and TB_VCT_2, postmitotic VCT clusters; TB_VCT_IFN, VCT subset exhibiting active interferon response.

For further analysis, we merged analogous subclusters into three major clusters— VCT, EVT, and SCT—and evaluated pathway activities in these aggregated clusters for PE versus controls (Figure 4). As before, hypoxia signaling was most pronounced in EVT, with activity highest in eoPE compared to controls. The SCT fraction in eoPE exhibited the next greatest hypoxia activity. Across all three populations, hypoxia pathway activity was elevated in PE relative to normal samples. Additionally, NFκB, PI3K, TGF-β, TNF-α, and TRAIL pathways were most active in EVT, consistent with our earlier observations.

**Fig. 4.**
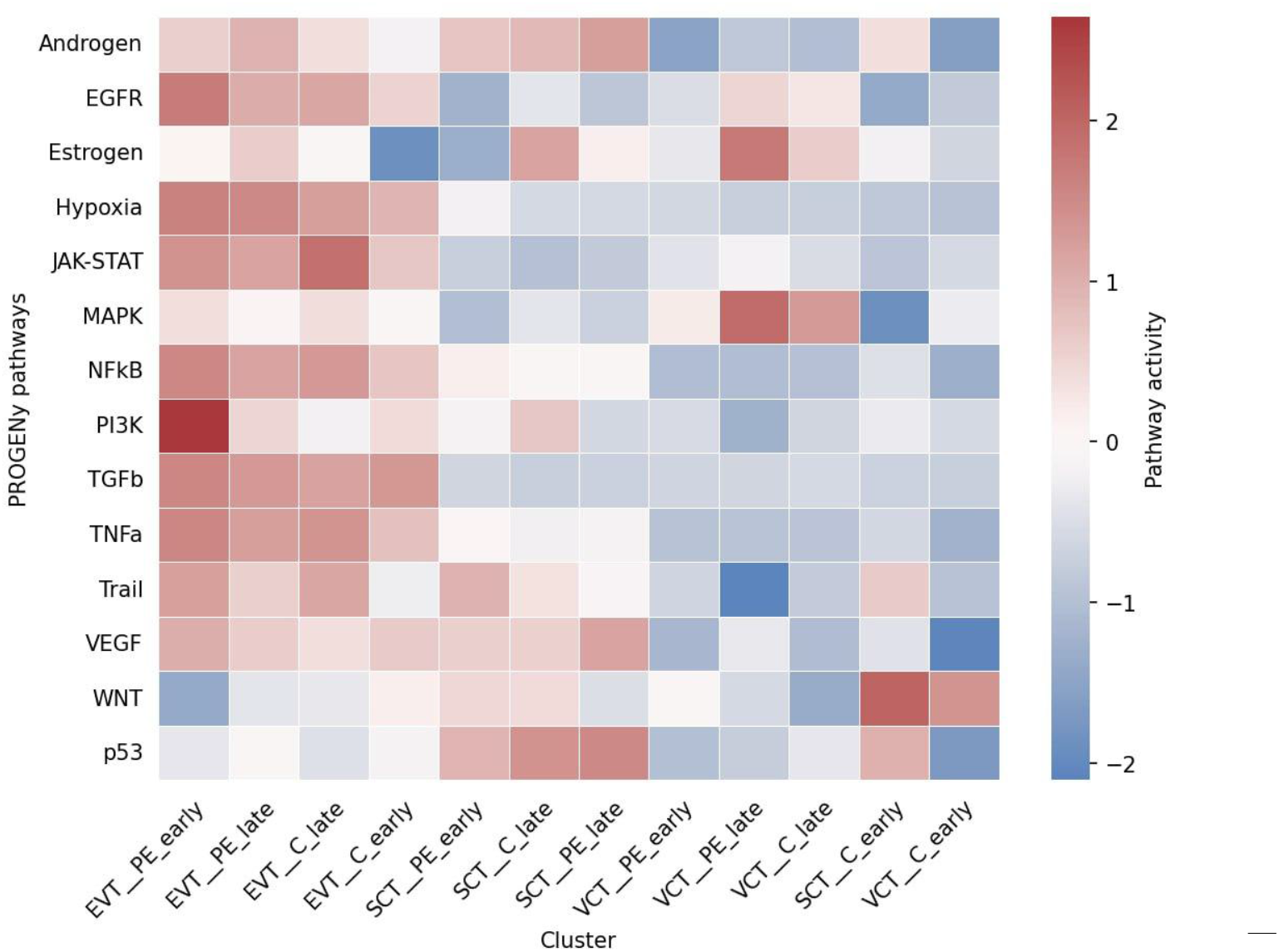
PROGENy-inferred signaling pathway activities in aggregated trophoblast clusters from scRNA-seq data [16]. EVT, extravillous trophoblast; SCT, syncytiotrophoblast; VCT, villous cytotrophoblast; PE_early, eoPE samples; C_early, controls for eoPE (delivery <34 weeks); PE_late, loPE samples; C_late, controls for loPE (delivery ≥34 weeks).

#### 3.1.3. Differential gene expression analysis identifies nine core hypoxia markers shared across all trophoblast clusters

Given the elevated hypoxia signaling in eoPE consistent with placental dysfunction, we concentrated on eoPE samples and matched controls. Differential expression analysis revealed the fewest significant changes in EVT, with 76 genes altered (38 upregulated and 38 downregulated in PE). In SCT, 458 genes were differentially expressed (241 upregulated, 217 downregulated), and in VCT, 368 genes (291 upregulated, 77 downregulated). Intersection analysis (Figure 5) identified nine genes— *EBI3, CST6, FN1, RFK, COL17A1, LDHA, PKP2, RPS4Y1*, and *RPS26*—that were consistently upregulated, and six genes—*CGA, PLCG2, LY6K, ATP1A4, MTRNR2L12*, and *MTRNR2L8*—that were consistently downregulated across all three trophoblast populations.

**Fig. 5.**
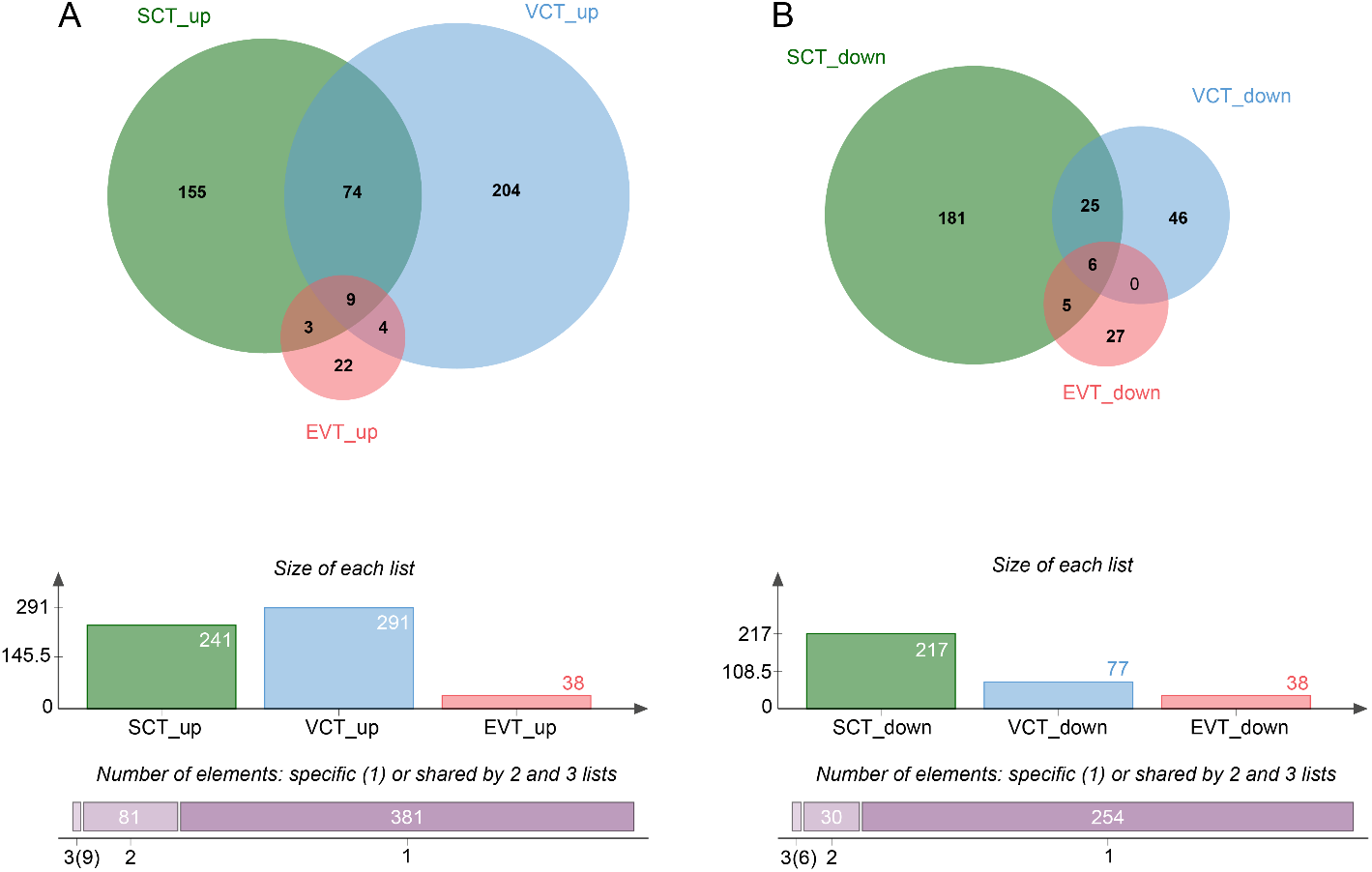
Differential gene-expression changes in trophoblast clusters. **A**. Upregulated genes. **B**. Downregulated genes. *Top*: Venn diagrams of gene-list intersections. *Middle*: number of differentially expressed genes per cluster. *Bottom*: counts of genes common to all three populations, shared by any two, or unique to each. EVT, extravillous trophoblast; SCT, syncytiotrophoblast; VCT, villous cytotrophoblast.

### 3.2. Analysis of BeWo b30 cell sequencing results

#### 3.2.1. Oxyquinoline derivative induces stronger hypoxia, TGF-β, TNF-α, and TRAIL signaling in BeWo b30 cells than CoCl2

We induced HIF-1α stabilization in BeWo b30 cells using cobalt(II) chloride and OD, the latter providing more selective PHD inhibition. PROGENy analysis (Figure 6) demonstrated that both agents elevated hypoxia pathway activity, with OD eliciting the greatest response. TGF-β, TNF-α, and TRAIL pathway activities were also higher following OD treatment, whereas NFκB activation was marginally greater with CoCl_2_. Notably, PI3K signaling increased under cobalt treatment but decreased with OD.

**Fig. 6.**
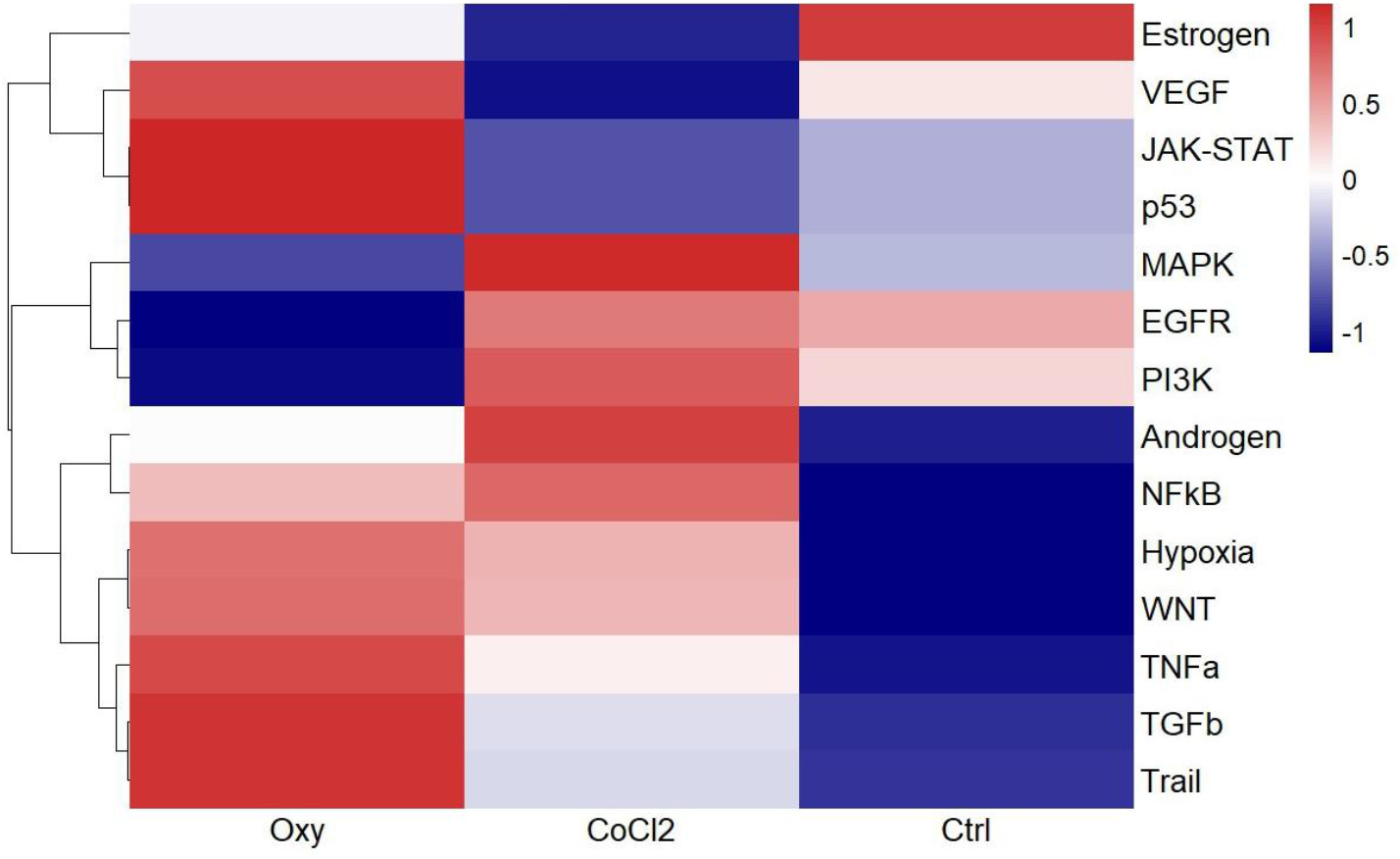
Activity of PROGENy-inferred signaling pathways following hypoxia induction in BeWo b30 cells. Oxy, cells treated with 5 µM oxyquinoline derivative for 24 h; CoCl_2_, cells treated with 300 µM cobalt(II) chloride for 24 h; Ctrl, untreated control cells in standard medium.

#### 3.2.2. Differential gene expression shows distinct and shared metabolic and mitochondrial pathway regulation upon chemical hypoxia induction

We quantified changes in protein-coding gene expression in BeWo b30 cells treated with OD or cobalt(II) chloride, defining significance as |FC| ≥ 2 at adjusted p < 0.05. OD exposure altered 1,284 genes (596 upregulated, 688 downregulated), whereas CoCl_2_ treatment affected 3,032 genes (1,397 upregulated, 1,635 downregulated). Intersection analysis (Figure 7) with annotation of common genes via DAVID [24] revealed that shared upregulated genes were enriched in ascorbate and iron metabolism (p = 0.004), transcriptional regulation (p = 0.009), autophagy (p = 0.010), circadian rhythms (p = 0.022), and the HIF-1 signaling pathway (p = 0.036). Shared downregulated genes clustered in mitochondrial function (p < 0.0001), proline biosynthesis (p = 0.003), cell division including cell-cycle regulation via ubiquitin-dependent proteolysis (p = 0.010), fatty-acid metabolism (p = 0.019), histone methylation (p = 0.021), kinesin-mediated intracellular transport (p = 0.032), and SLC25-mediated mitochondrial transport (p = 0.041).

**Fig. 7.**
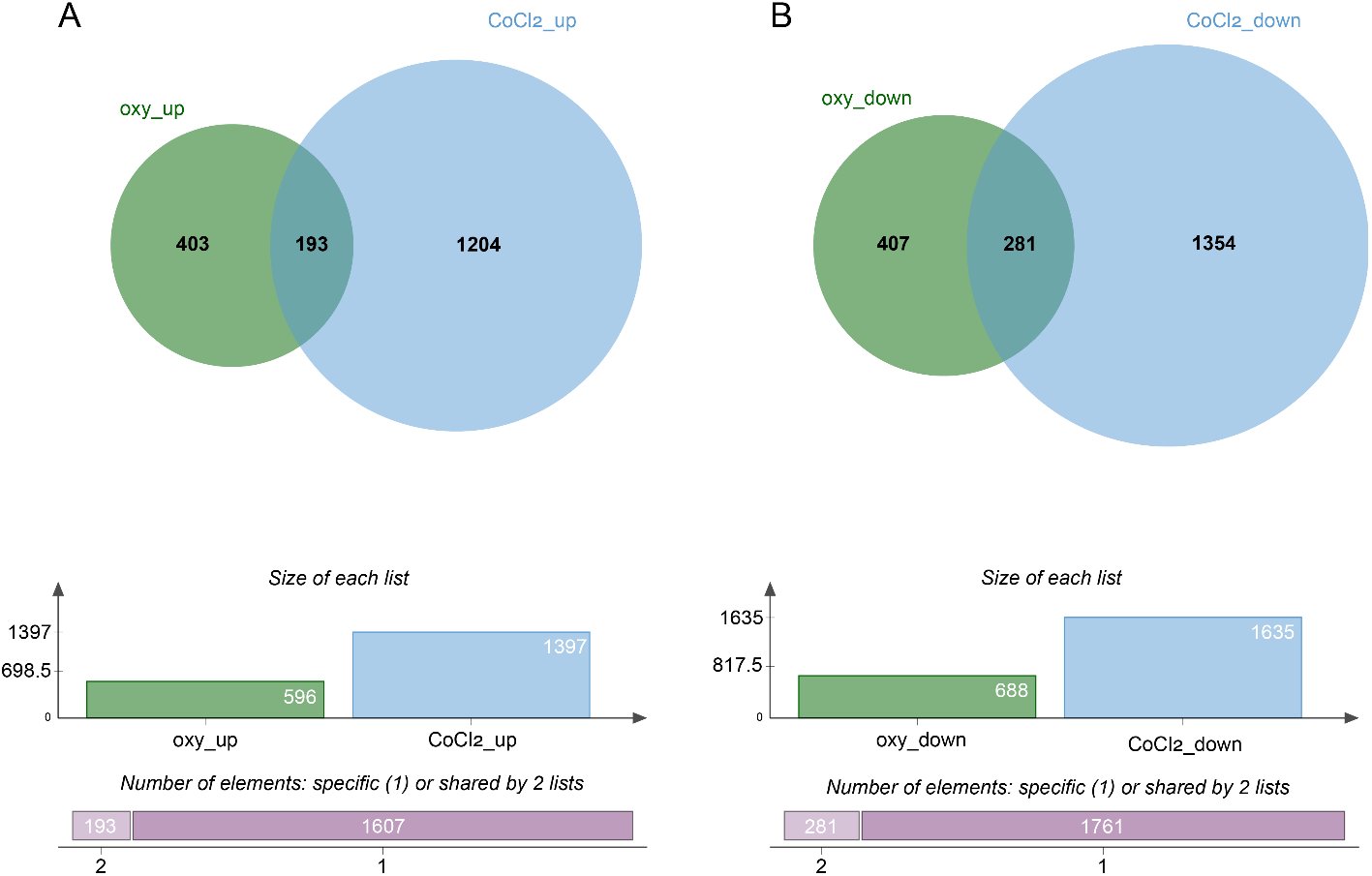
Differential gene-expression changes in BeWo b30 cells following chemical hypoxia induction. **A**. Upregulated genes. **B**. Downregulated genes. *Top*: Venn diagrams of gene-list intersections. *Middle*: number of differentially expressed genes per condition. *Bottom*: counts of genes shared by two conditions or unique to each. CoCl_2_, cobalt(II) chloride; oxy, oxyquinoline derivative.

#### 3.2.3. Oxyquinoline derivative faithfully reproduces EBI3 and COL17A1 upregulation observed in preeclamptic trophoblasts

We next assessed whether genes upregulated across all trophoblast clusters in eoPE were similarly regulated in BeWo b30 cells treated with CoCl_2_ or OD (Figure 8). Notably, *EBI3, FN1*, and *COL17A1* exhibited inverse regulation upon CoCl_2_ exposure compared with PE trophoblasts—significantly so for *COL17A1*—whereas OD treatment recapitulated the direction of change seen in PE for all three genes. *RFK* expression remained unchanged with either hypoxia mimetic, and *PKP2* showed a nonsignificant opposite trend relative to PE. We also examined *HTRA4* and *FSTL3*—previously reported as overexpressed in extracellular vesicles from preeclamptic placentas [25]. *HTRA4* was overexpressed in SCT and VCT but downregulated in both treatments in BeWo b30 cells. *FSTL3* was overexpressed in VCT, tended to be overexpressed in EVT and SCT, was strongly induced by OD, yet repressed by CoCl_2_ in BeWo b30 (Figure 8).

**Fig. 8.**
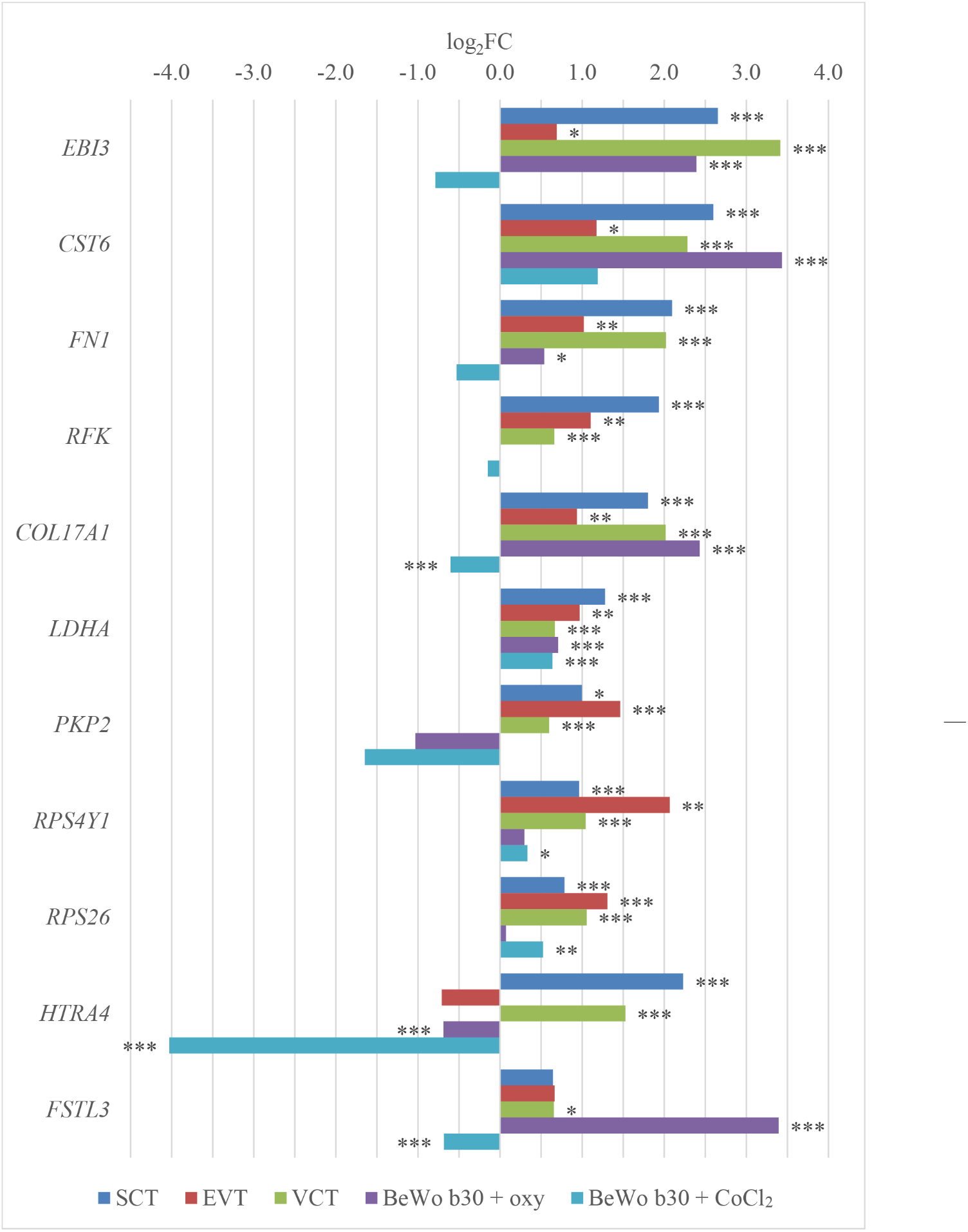
RNA-seq expression changes of key genes in preeclamptic trophoblast populations and following hypoxia induction in BeWo b30 cells. CoCl_2_, cobalt(II) chloride; oxy, oxyquinoline derivative; VCT, villous cytotrophoblast; EVT, extravillous trophoblast; SCT, syncytiotrophoblast. * p<0.05; ** p<0.01; *** p<0.001.

We next measured *EBI3, COL17A1, HTRA4, FSTL3*, and *CST6* expression by qPCR (Figure 9). *HTRA4* was strongly downregulated by CoCl_2_ and unchanged by OD. *FSTL3* followed the RNA-seq trend but did not reach statistical significance. *CST6* was significantly upregulated by both CoCl_2_ and OD. *EBI3* and *COL17A1* were significantly induced by OD, but showed non-significant changes in response to CoCl_2_.

**Fig. 9.**
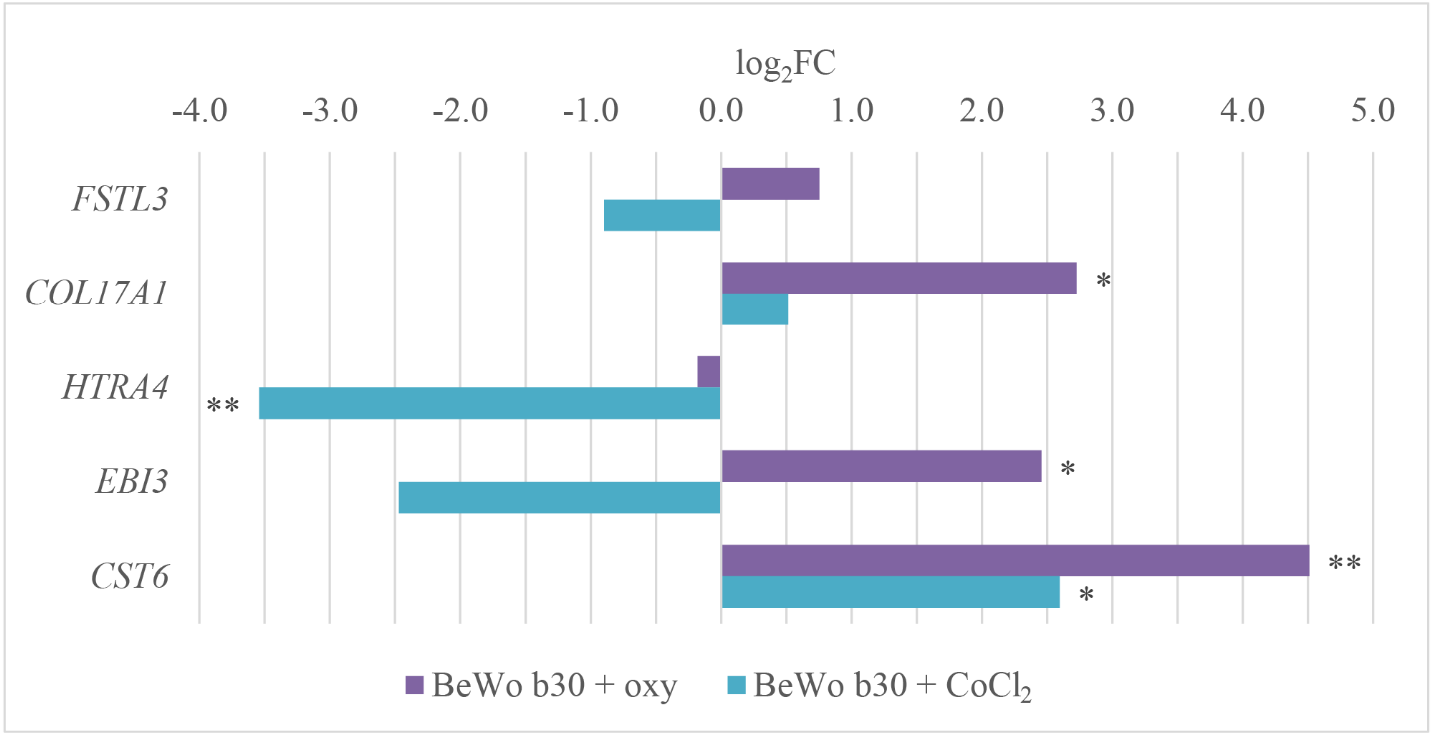
qPCR expression changes of selected genes following hypoxia induction in BeWo b30 cells. CoCl_2_, cobalt(II) chloride; oxy, oxyquinoline derivative. * p<0.05; ** p<0.01.

#### 3.2.4. Chemical hypoxia induction reveals hsa-miR-210-3p, hsa-miR-27a-5p, hsa-miR-193b-5p as shared PE markers and novel isomiRs

To pinpoint miRNAs linked to both PE placental pathology and trophoblast hypoxia, we referenced Awoyemi et al. (2024) [26], who reported 23 miRNAs significantly upregulated in placental tissue and extracellular vesicles from PE cases. Having performed small RNA sequencing on BeWo b30 cells treated with CoCl_2_ or OD, we searched our datasets for these miRNAs. Both treatments increased hsa-miR-210-3p, hsa-miR-27a-5p, and hsa-miR-193b-5p levels, while OD alone also elevated hsa-miR-455-3p (Figure 10).

**Fig. 10.**
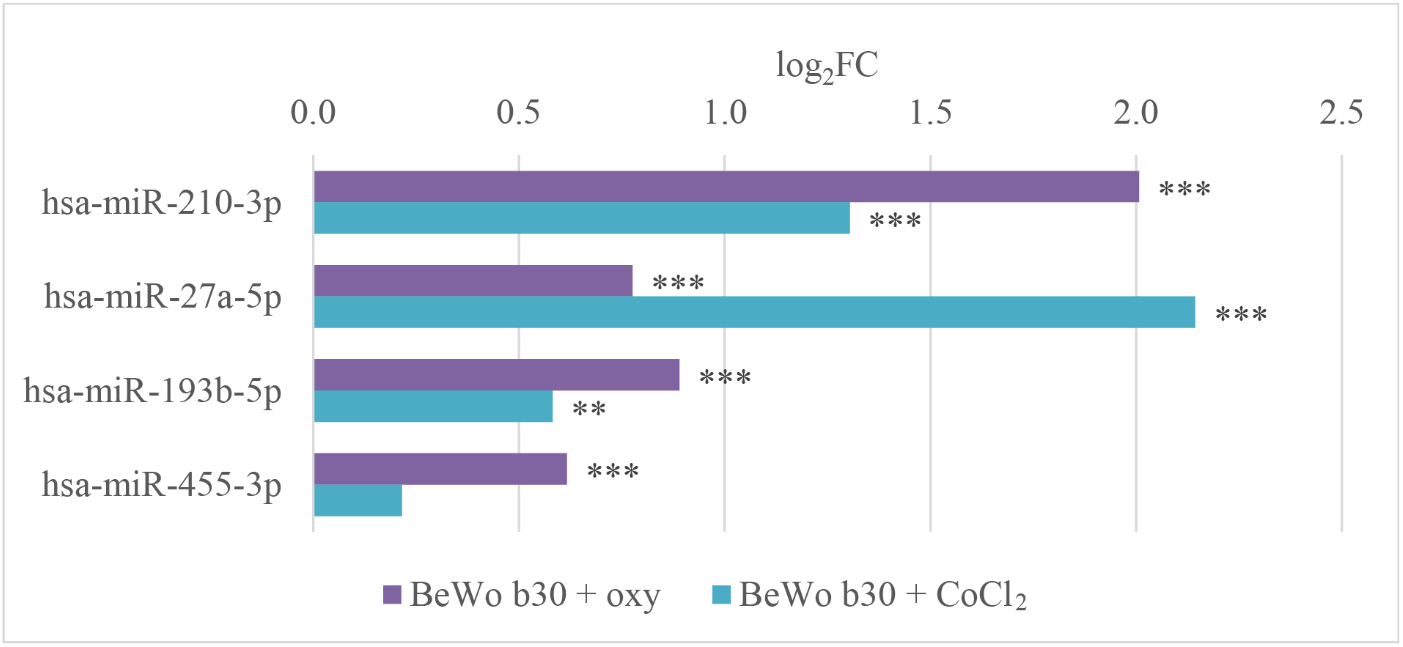
Expression changes of key microRNAs in BeWo b30 cells following hypoxia induction. CoCl_2_, cobalt(II) chloride; oxy, oxyquinoline derivative. ** p<0.01; *** p<0.001.

We also examined microRNA isoforms (isomiRs) in our small RNA-seq data. In addition to the canonical miRNAs, we observed significant upregulation of hsa-miR-9-5p isomiRs upon CoCl_2_ treatment. Unlike canonical sequence (5′-UCUUUGGUUAUCUAGCUGUAUGA-3′), the hsa-miR-9-5p|0|-3(+2G) isoform (5′-UCUUUGGUUAUCUAGCUGUAGG-3′) showed a 2.5-fold increase (p < 0.001), and the hsa-miR-9-5p|0|-3(+1G) isoform (5′-UCUUUGGUUAUCUAGCUGUAG-3′) increased 1.7-fold (p = 0.001). The latter isoform also rose 1.6-fold (p = 0.002) with OD treatment. Furthermore, OD exposure elevated the hsa-miR-92b-3p|0|+1 isomiR— extended by one uracil at the 3′ end—by 1.5-fold (p = 0.0001).

## 4. Discussion

Our integrated analysis demonstrates that eoPE is characterized by pronounced hypoxia signaling across all trophoblast subpopulations paralleling the heightened HIF-1α stabilization achieved with chemical hypoxia mimetics in BeWo b30 cells. By directly comparing PE placental scRNA-seq profiles with chemical hypoxia models, we reveal that oxyquinoline derivative (OD) more faithfully reproduces the direction and magnitude of PE-associated transcriptional and miRNA changes than CoCl_2_. These findings establish OD treatment of BeWo b30 cells as a superior *in vitro* paradigm for dissecting trophoblast hypoxic responses and uncover candidate mRNA and isomiR markers linked to placental dysfunction in PE.

Our single-cell RNA-seq analysis of placental trophoblasts in PE underscores both biological insights and key technical constraints. Notably, the VCT:SCT ratio was markedly skewed—3.8:1 in loPE and 5:1 in eoPE—far from physiological proportions, highlighting an inherent limitation of scRNA-seq for placental tissue [15]. Because the SCT is a large multinucleated syncytium, it is largely lost during tissue dissociation and the microfluidic cell-size filtration required for the 10x Genomics platform, leading to substantial underestimation of SCT abundance. This bias can misrepresent trophoblast composition and thus hinder accurate modeling of PE pathophysiology. Single-nucleus RNA-seq (snRNA-seq) offers a promising alternative—capturing syncytial nuclei at approximately 50-fold higher efficiency than scRNA-seq [27,28]—but introduces trade-offs, including reduced sensitivity for mature transcripts and potential cross-contamination of nuclear RNA from neighboring cell types.

The PROGENy-based signaling pathway analysis uncovers fundamental mechanisms driving PE pathogenesis and distinguishes early-onset from late-onset disease. By inferring activity of 14 core oncogenic and signaling cascades from their downstream target genes, PROGENy offers a more robust functional readout than single-gene analyses [18]. This integrative framework is especially valuable for trophoblast studies, integrating complex transcriptional shifts into coherent activity signatures. Moreover, PROGENy implicitly captures post-translational and regulatory effects that may not manifest at the mRNA level [18]. Although originally designed for bulk RNA-seq, PROGENy has demonstrated high applicability and performance in single-cell contexts [29].

The analysis revealed pronounced activation of hypoxia pathways in PE, with a clear gradient of activity (eoPE > loPE > controls) that aligns with current models of PE pathophysiology and underscores the central role of placental hypoxia in syndrome development [30]. Notably, hypoxia signaling peaked in EVT and the SCT_PSG subset with elevated placenta-specific genes. This pattern is biologically coherent: EVT migrate into the decidua, distant from maternal vasculature, creating a naturally hypoxic microenvironment [31]. Elevated hypoxia pathway activity in SCT_PSG likely reflects the metabolic specialization of this fraction—the most viable SCT population retained after dissociation-induced losses in scRNA-seq [16].

The coordinated activation of NFκB, PI3K, TGF-β, TNF-α, and TRAIL pathways in eoPE highlights a complex interplay of inflammatory and apoptotic processes characteristic of severe placental dysfunction [4]. NFκB, a master regulator of inflammation, drives proinflammatory cytokine expression and contributes to trophoblast invasion during placentation [32]. Concurrent PI3K activation likely represents a cell-survival response to hypoxic stress [33]. TGF-β signaling, implicated in tissue remodeling, may both promote placental fibrosis and modulate immune tolerance in PE [34]. Notably, TRAIL (TNF-related apoptosis-inducing ligand) is secreted by trophoblasts to induce apoptosis of vascular smooth muscle cells during spiral artery remodeling; however, its overactivation in PE may exacerbate trophoblast loss and impair placentation [35]. The predominance of these pathway activities in EVT underscores their pivotal role as invasive cells orchestrating uteroplacental blood flow remodeling [31].

Chemical induction of HIF signaling in BeWo b30 cells with CoCl_2_ and the oxyquinoline derivative (OD) recapitulated key features of placental hypoxia and revealed distinct signaling responses. Both agents robustly activated the hypoxia pathway, but OD—via selective inhibition of HIF prolyl hydroxylases—elicited a markedly stronger response than CoCl_2_. OD also drove greater activation of TGF-β, TNF-α, and TRAIL pathways, reflecting intensified inflammation, apoptosis, and matrix remodeling under severe hypoxic stress without confounding ionic effects [34–36]. In contrast, CoCl_2_ induced more pronounced NF-κB activation, likely due to cobalt-ion– mediated oxidative stress and direct modulation of inflammatory signaling [32,37].

The differential expression analysis in eoPE trophoblasts revealed both shared and lineage-specific adaptations to hypoxic and inflammatory stress. EVT exhibited the fewest changes (76 genes), consistent with their relative resilience to placental stress and their underrepresentation in scRNA-seq. Intersection of upregulated genes across SCT, VCT, and EVT identified nine core responders: *EBI3, CST6, FN1, RFK, COL17A1, LDHA, PKP2, RPS4Y1*, and *RPS26*, likely reflecting a unified trophoblast response to PE and hypoxia. Notably, placental perfusion studies have reported elevated *COL17A1* and *EBI3* in small and medium/large extracellular vesicles from PE placentas, suggesting trophoblast secretion into maternal circulation [25]. Those vesicles also carried increased *FLNB, HTRA4*, and *PAPPA2*—genes we found overexpressed in SCT and VCT—and *INHBA*, which was similarly elevated in SCT in our scRNA-seq data.

*EBI3* encodes a subunit of the immunomodulatory cytokines IL-27 and IL-35. In our BeWo b30 hypoxia model, CoCl_2_ tended to downregulate *EBI3* expression, whereas OD induced a 5.5-fold upregulation, suggesting it more faithfully mimics PE-associated trophoblast hypoxia. In normal pregnancy, IL-27–associated EBI3 is highly expressed in SCT and EVT across all trimesters but is low in VCT [38], and IL-35–associated EBI3 is present in first-trimester primary trophoblasts and HTR-8 cells [39]. Decidual placental tissue from PE cases shows elevated *EBI3* mRNA compared with normotensive controls [40], and circulating EBI3 is increased in eoPE but not loPE [41,42]. Extracellular vesicles from PE placentas also exhibit higher EBI3 mRNA in small vesicles and a trend toward increase in medium/large vesicles [25]. Metformin reduces lncRNA H19 in trophoblast and endothelial cells, thereby increasing miR-216-3p and suppressing *EBI3*, hinting at a therapeutic mechanism in PE [43]. To our knowledge, no previous trophoblast cell-line model demonstrated hypoxia-induced *EBI3* upregulation in a PE context; however, STOX1 overexpression in JEG-3 cells produced a 12-fold *EBI3* increase [44].

All three trophoblast fractions in PE exhibited upregulated *COL17A1*, which encodes the α1 chain of type XVII collagen. Unlike fibrillar collagens, type XVII is a transmembrane protein whose ectodomain (LAD-1) is released by proteolytic cleavage [45]. In our BeWo b30 hypoxia model, OD treatment induced a 6.6-fold increase in *COL17A1* expression, whereas CoCl_2_ failed to alter its expression. Prior studies have reported elevated *COL17A1* in PE placentas [46–50], correlating with increased immune cell infiltration [47].

The *FN1* gene encodes fibronectin, which was upregulated across all trophoblast fractions in PE. In BeWo b30 cells, OD induced a 1.5-fold increase in *FN1* expression, whereas CoCl_2_ caused a 1.5-fold decrease, underscoring divergent modeling of hypoxic responses by these inducers. Circulating FN1 is elevated in PE patients’ plasma, particularly in severe, antihypertensive-resistant cases [50–52]. Placental FN1 overexpression may be driven by hypoxia-induced histone lactylation via lactate production [53]. Corroborating this, we observed *LDHA* upregulation in all trophoblast fractions in eoPE—and a 1.6-fold increase in *LDHA* in BeWo b30 under both treatments (adj. p < 0.0001)—indicating enhanced glycolytic flux. In PE placentas, *FN1, ITGA5*, and *FOS* are hypomethylated and overexpressed [54]. Consistently, *ITGA5* expression rose 3.9-fold in SCT and 1.5-fold in EVT in eoPE (adj. p = 0.004 and 0.025, respectively), with no significant change in loPE. OD and CoCl_2_ treatments increased *ITGA5* by 1.6- and 1.5-fold in BeWo b30 (adj. p < 0.0001). Integrin A5 belongs to the family of RGD receptors [55]. We previously demonstrated *ITGA5* upregulation in inflammatory bowel disease tissues and in Caco-2 cells under hypoxia [56], suggesting a broader role for ITGA5 in conditions characterized by hypoxic and immune dysregulation, such as PE and inflammatory bowel disease.

*CST6* encodes cystatin E/M, a cysteine protease inhibitor. Its expression is elevated in PE placental tissue and maternal serum [50,57,58], and in syncytiotrophoblasts under hypoxia (1% vs. 8% O_2_) [57]. In BeWo b30 cells, both hypoxia inducers increased *CST6* expression. However, unlike other markers, CST6 has not been reported in PE-associated extracellular vesicles, potentially limiting its utility as a circulating biomarker.

Analysis of microRNA expression in BeWo b30 cells following chemical hypoxia induction uncovered both established PE markers and novel isomiRs that may exert additional regulatory functions. Three miRNAs—hsa-miR-210-3p, hsa-miR-27a-5p, and hsa-miR-193b-5p—were significantly upregulated by both CoCl_2_ and OD. hsa-miR-210-3p, a prototypical “hypoxamiR,” is HIF-1α–dependent and modulates metabolism, angiogenesis, and cell survival under low-oxygen conditions [59]. Elevated hsa-miR-27a-5p and hsa-miR-193b-5p have been reported in placental exosomes from PE patients [26]. Although miR-27a’s role in PE remains underexplored, we previously observed increases in hsa-miR-27a-3p and -5p in hypoxic BeWo b30 cells and implicate them in TGF-β signaling regulation in PE macrophages [60]. Circulating miR-27a is also elevated in PE and can inhibit trophoblast migration via SMAD2 targeting [61,62]. hsa-miR-193b-5p has likewise been implicated in PE pathogenesis [63]. OD treatment specifically increased hsa-miR-455-3p, previously found elevated in PE placental tissue and exosomes associated with gestational diabetes [64,65]. Beyond canonical miRNAs, we detected significant upregulation of hsa-miR-9-5p and hsa-miR-92b-3p isomiRs, which differ by 3′-end nucleotide additions or substitutions without predicted target shifts.

In summary, our study provides the first direct comparison of *in vivo* trophoblast hypoxic responses in eoPE placentas with *in vitro* chemical HIF stabilization in BeWo b30 cells, revealing that OD treatment more accurately mirrors the transcriptional, pathway-activity, and miRNA signatures of PE than CoCl_2_. We identify a core set of mRNA and miRNA markers (including *EBI3, COL17A1*, hsa-miR-27a-5p, and hsa-miR-193b-5p) and novel isomiRs (notably hsa-miR-9-5p and hsa-miR-92b-3p variants) that delineate trophoblast adaptation to hypoxic and inflammatory stress. These molecular signatures offer promising candidates for future biomarker validation and mechanistic studies of placental dysfunction in PE.

## Declaration of competing interest

None.

## Funding

The study for the sections concerning analysis of single-cell sequencing data was performed within the framework of the Basic Research Program at HSE University. The study for the sections concerning BeWo b30 cell model was funded by the Russian Science Foundation grant No. 24-14-00382, https://rscf.ru/en/project/24-14-00382/

## Declaration of generative AI and AI-assisted technologies in the writing process

During the preparation of this work the authors used Perplexity AI tool in order to improve the readability and language of the manuscript. After using this tool, the authors reviewed and edited the content as needed and take full responsibility for the content of the published

## Notes

https://www.ncbi.nlm.nih.gov/geo/query/acc.cgi?acc=GSE308908

### Competing Interest Statement

The authors have declared no competing interest.

